# Molecular determinants of Yellow Fever Virus pathogenicity in Syrian Golden Hamsters: one mutation away from virulence

**DOI:** 10.1101/249383

**Authors:** Raphaëlle Klitting, Laura Roth, Félix A. Rey, Xavier de Lamballerie

## Abstract

Yellow fever virus (*Flavivirus* genus) is an arthropod-borne pathogen which can infect humans, causing a severe viscerotropic disease with a high mortality rate. Adapted viral strains allow the reproduction of yellow fever disease in hamsters with features similar to the human disease. Here, we used the Infectious Subgenomic Amplicons reverse genetics method to produce an equivalent to the hamster-virulent strain, Yellow Fever *Ap7*, by introducing a set of 4 synonymous and 6 non-synonymous mutations into a single subgenomic amplicon, derived from the sequence of the *Asibi* strain. The resulting strain, Yellow Fever *Ap7M*, induced a disease similar to that described for *Ap7* in terms of symptoms, weight evolution, viral loads in the liver and lethality. Using the same methodology, we produced mutant strains derived from either *Ap7M* or *Asibi* viruses and investigated the role of each of *Ap7M* non-synonymous mutations in its *in vivo* phenotype. This allowed identifying key components of the virulence mechanism in hamsters. In *Ap7M* virus, the reversion of either E/Q27H or E/D155A mutations, led to an important reduction of both virulence and *in vivo* replicative fitness. In addition, the introduction of the single D155A *Ap7M* mutation within the E protein of the *Asibi* virus was sufficient to drastically modify its phenotype in hamsters towards both a greater replication efficiency and virulence. Finally, inspection of the *Asibi* strain E protein structure combined to *in vivo* testing revealed the importance of an exposed α-helix in domain I, containing residues 154 and 155, for *Ap7M* virulence in hamsters.

## INTRODUCTION

The genus *Flavivirus* (family, *Flaviviridae*) includes more than 60 enveloped, positive and single-stranded RNA viruses. It was named after the jaundice characteristic of the liver dysfunction caused by the infection with Yellow fever virus (YFV), the funding member of the genus (1). Like a majority of flaviviruses, YFV is an arthropod-borne human pathogen (arbovirus). It was one of the first viruses of humans to be identified, isolated (2), propagated *in vitro* (3-5) and studied by genomic sequencing (6, 7). The YF virion consists in a spherical, enveloped particle of 50nm in diameter (8). The capped, 11 kilobases genome includes a single open reading frame (ORF) flanked at its 5’ and 3’ termini by structured, non-coding regions, necessary to viral RNA translation and replication (9). ORF translation gives rise to a polyprotein that is cleaved co- and post-translationally into 3 structural and 7 non-structural proteins. The 3 structural proteins are the constituents of the capsid (C), the envelope (E) and the membrane (for which both an immature (prM) and a mature (M) form have been described) (1). The E protein is present as a homodimer at the surface of the viral envelope. Each E monomer includes a central domain (I), a dimerization domain (II) and a putative receptor-binding domain (III) (10). The E protein is involved in receptor binding and membrane fusion (11-13). Non-synonymous mutations in this protein have been reported to impact viral tropism and virulence (14-18).

YFV is endemic in the tropical regions of Africa and South-America, where its natural upkeep is conditioned by the presence of both its mosquito vector(s) and its primate host(s). In Africa, YFV is maintained through a transmission cycle between sylvatic vectors (several species (sp.) including *Aedes (Ae.) africanus* and *Ae. furcifer* and non-human primate (NHP) hosts (genera *Cerecopithecus, Colubus*, and *Erythrocebus*) (sylvatic cycle) (19, 20). Humans get infected when bitten by sylvatic mosquitoes that previously fed on viremic monkeys. Inter-human transmission can occur when infected and naïve human hosts are confined together with a competent bridge or domestic vector (notably *Ae. aegypti* and *Ae. bromeliae* sp.) (sylvatic/savanna and urban cycles) (21). Lastly, transovarial transmission in the mosquito vector may also contribute to the spread of the virus (6). YFV circulation was documented in numerous regions in South America (notably in Brazil, Argentina, Paraguay, Bolivia and Peru) and involves mosquito species from the genera *Haemagogus* and *Sabethes* that ensure transmission amongst NHPs and from NHPs to humans (22). YFV infection was notably described in spider monkeys *(Ateles* sp.), owl monkeys *(Aotus* sp.) and howler monkeys from the genera *Alouatta*, for which important lethality was recorded, reaching degrees of severity that were never reported in monkeys from the African continent (23, 24).

In humans, YFV causes a severe viscerotropic disease, infecting primarily the liver in association with high fever. The virus can affect other tissues such as spleen, kidney, lymph nodes or heart (6, 20). The severity of YF infections ranges from “inapparent” cases *(i.e*. that do not call medical attention) to fatal diseases with a ratio of inapparent to apparent infection of approximately 7–12:1 (estimated from field studies (25, 26)) and a mortality rate between 20 and 50% amongst symptomatic cases (27).

To date, no antiviral drug is available for YFV treatment, which remains limited to symptomatic and supportive care. Although different molecules have shown *in vitro* and/or *in vivo* activity against YFV, none is available for clinical use (6, 27). An effective live-attenuated vaccine (strain 17D) was developed in 1936 by M. Theiler. It served as early as 1938 in Brazil (28, 29) and two of its substrains (17DD and 17D-204) are still widely used for vaccine manufacturing (30), with an annual distribution of 20 to 60 million doses (6).

YFV species includes one serotype that groups together 5 African and 2 South-American genotypes (31, 32). The virus originates in Africa and crossed the Atlantic Ocean to the Americas during the slave trade. Since then, large epidemics have been described in these regions. Human activities ensure further YF dissemination from endemic areas to Europe, North America and the Caribbean (6). Despite vector-control plans and although 500 million doses of vaccine were distributed over the last 50 years (6), YFV is still causing important outbreaks as recently reported in Africa (33) and in South America (34). This is mainly explained by the efficient maintenance of the virus through its sylvatic cycle from which it can start interhuman transmission cycles. With around 900 million people at risk in endemic areas (26), there is still a need for further understanding YFV molecular mechanisms of replication and virulence in order to identify new therapeutic targets and to develop better countermeasures during outbreaks (34).

Several animal models allow to reproduce some aspects of the clinical disease caused by YFV in humans (35). NHPs (notably sp. *Macaca mulatta*) reproduce well the features of human YF but the use of such models raises ethical, regulatory and biosafety concerns. Transgenic mice models with defects in the interferon signaling pathway are easier to set up but they poorly reproduce the disease observed in humans. YF infection in hamsters (sp. *Mesocricetus Auratus*) can cause a viscerotropic disease that affects the liver with symptoms similar to those of the human disease. This model requires the use of adapted strains (15, 35-37) such as the pathogenic YF hamster-viscerotropic strain *Asibi*/hamster p7 (*Ap7*), which was described in 2003 (15). It was derived from the strain *Asibi*/hamster p0 (*Ap0*), avirulent in hamsters. After 7 passages in hamster, the derived strain *Ap7* induced a symptomatic liver infection along with a robust viremia and a high mortality rate. Sequencing of both the parental and daughter strains allowed the identification of 14 nucleotide changes amongst which 7 caused amino-acid modifications. A majority of those mutations fell within the E protein, suggesting an important role of this protein for the observed virulence in hamsters.

In this work, a subset (10) of the mutations described in YF *Ap7* virus was introduced into the genome of the *Asibi* strain using the Infectious Subgenomic Amplicons (ISA) reverse-genetics method (38). The resulting virus, YF Asibi/hamster p7 Marseille (*Ap7M*), was inoculated to hamsters and the hamster-lethal infection described for YFV *Ap7* strain was reproduced. Each of the non-synonymous mutations included in strain *Ap7M* was then separately reverted and finally, some of the amino acids responsible for the hamster-virulent phenotype of *Ap7M* virus were identified by introducing single mutations into the genome of the *Asibi* strain.

## MATERIALS AND METHODS

*Protocols regarding cell culture (virus stock production and virus titration (TCID50)), PCR amplification and fusion (subgenomic amplicon production), transfection and virus stock production, quantitative real-time PCR (qRT-PCR) assays, sample collection as well as next-generation sequencing are detailed in the supplementary materials*.

### Cells and animals

Viruses were produced and titrated in Baby hamster kidney BHK21 cells (ATCC, number CCL10). *In vivo* infection was performed in three-week-old female Syrian Golden hamsters (*Mesocricetus Auratus*, Janvier laboratories).

### Ethics statement

Animal protocols were reviewed and approved by the ethics committee “Comité d’éthique en expérimentation animale de Marseille—C2EA—14” (protocol number 2015042015136843-V3#528). All animal experiments were performed in compliance with French national guidelines and in accordance with the European legislation covering the use of animals for scientific purposes (Directive 210/63/EU).

### In-silico *Asibi* mutant strains design

All mutant strains used in this study were produced using the *Infectious Subgenomic Amplicons* (ISA) method (38) by which infectious virus can be recovered after transfection into permissive cells of overlapping subgenomic DNA fragments covering the entire genome.

#### Ap7-derived hamster-adapted strains

The *Ap7M* hamster-adapted strain was created by introducing 10 mutations in the 1^st^ subgenomic fragment (FI) used for virus production with the ISA method (see Fig 1). FI region involves nucleotide genome positions 1-3919 that code for proteins C to NS2A. Amongst the 14 changes described in strain *Ap7* (9), 6 non-synonymous (NS) and 4 synonymous (S) mutations were introduced into the genome of the YFV strain *Asibi* (34) (Genbank accession number (AN): AY640589). NS changes were located in the E (Q27H, D28G, D155A, K323R, K331R) and NS2A (T48A) proteins. We could not identify the synonymous mutation described in the NS1 coding region (theoretically in position 3274 on the complete coding sequence (CDS)). In position 4864 (NS3 coding region), our reference strain was similar to strain *Ap7* and not to the original Asibi/hamster p0 (Ap0) strain described by McArthur (15). Hence, aside from three mutations (1 NS, 2 S) strain *Ap7M* is similar to *Ap7* strain. Details regarding the genomic composition of strains *Ap0, Ap7* and *Ap7M* are available in Fig 2.

**Figure 1.**
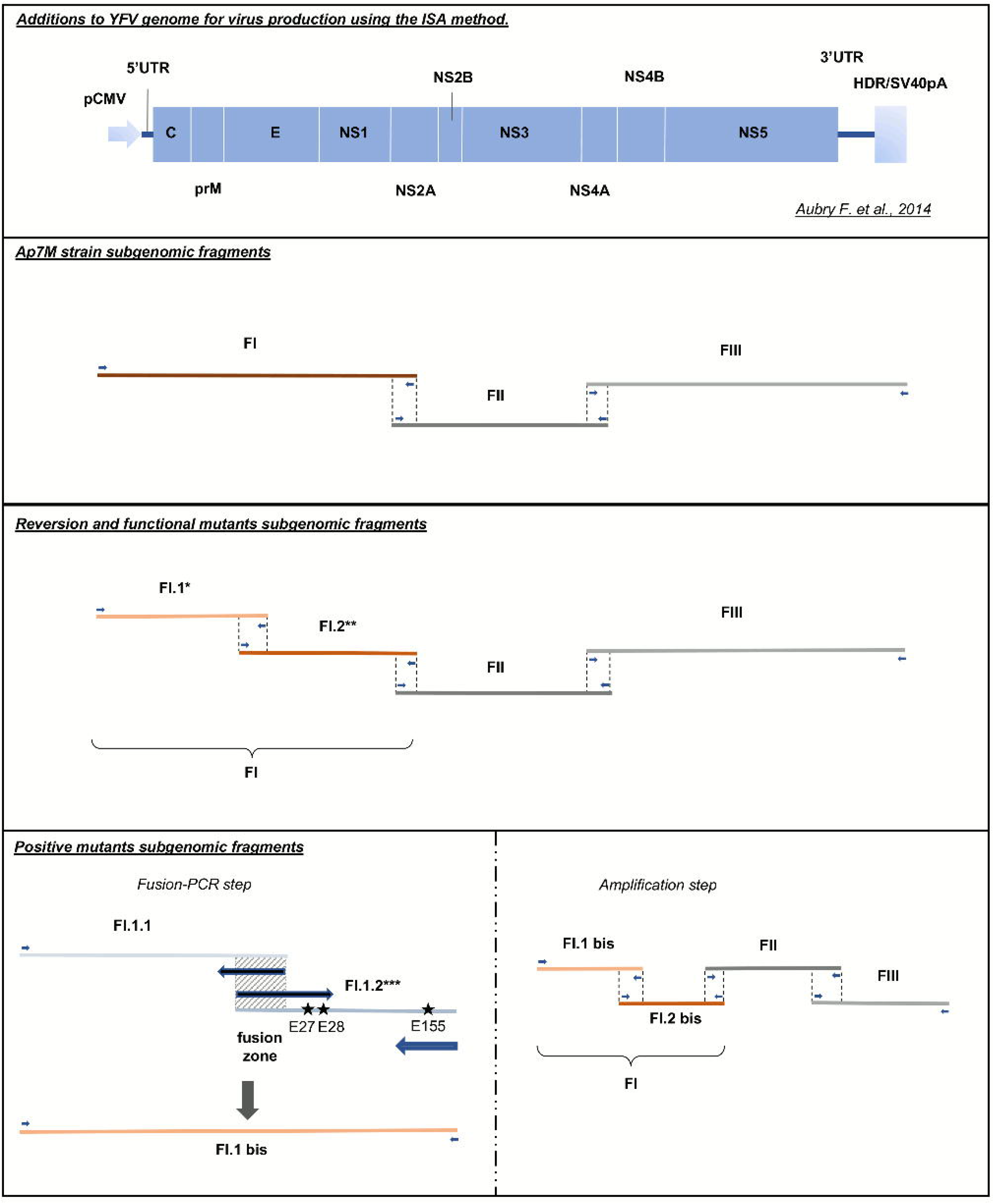
Viral strains production using the Infectious Subgenomic Amplicons (ISA) method. All viral strains were produced using the ISA method. Transfection schemes are detailed for all viruses. * For each of the envelope (E) reversion and functional mutants, a different FI.1 fragment was amplified from the corresponding plasmid. ** For YFV *Ap7M* NS2A/A48T reversion mutant production, FI.1 and FI.2 fragments were amplified from *Ap7M* and *Asibi* FI plasmids, respectively. *** For each of the E positive mutants, E/Q27H, E/D28G and E/D155A mutations were introduced in fragment FI.1.2 separately or as a combination of 2 to 3 changes, using modified primers (see supplementary Table S1). FI.1.1 and FI.1.2 fragments were subsequently fused in fragment FI.1 bis (PCR-fusion step).

**Figure 2.**
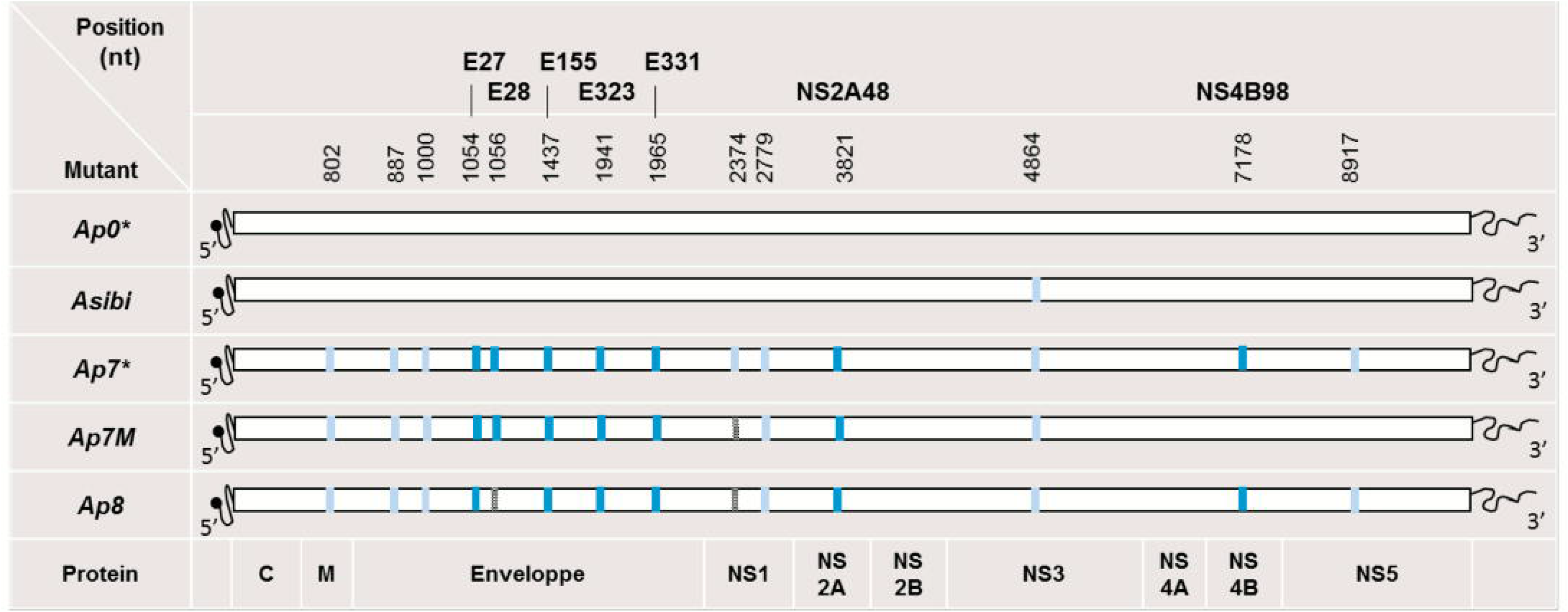
Description of ApO, Asibi, Ap7, Ap7M and Ap8 strains. All viral strains produced and used in this study are detailed above. Synonymous mutations are indicated by light blue rectangles and non-synonymous mutations by turquoise blue rectangles. Non-synonymous mutations are described in bold characters with a letter indicating the protein affected and the position within the protein sequence. Within the polyprotein sequence, the mature E, NS2A and NS4B protein sequences start at amino acid (AA) positions 286, 1131 and 2257, respectively. *\*Ap0* and *Ap7* strains description was obtained from Mc Arthur *et al*. 2003. *Asibi* strain sequence was downloaded from Genbank (AN: AY640589). Absent mutations are indicated by grey rectangles. We were not able to identify the synonymous mutation on the complete coding sequence (CDS) (grey rectangles). E/D28G mutation was not found in strain *Ap8* sequence.

#### Ap7M-derived reversion mutant strains

For each of the 6 non-synonymous changes introduced in the *Ap7M* sequence, a “reversion” mutant was created by reverting the mutation back to the *Asibi* strain sequence, leaving other mutations unchanged.

#### Asibi-derived positive mutant strains

“Positive” mutant strains were designed by selectively introducing mutations E/Q27H, E/D28G and E/D155A alone or in combination into the sequence of the wild-type strain *Asibi*.

#### Ap7M-derived functional mutant strains

“Functional” mutants were designed by replacing either Q27H or D155A changes by a hypothetically functionally equivalent NS mutation. For creating the E/T154A mutant, the D155A change was replaced by a T154A mutation (*i.e*. by a mutation on the adjacent amino acid residue within the E protein). For creating the E/Q27A and E/Q27N functional mutants, the Q27H change was replaced by Q27A and Q27N mutations, respectively. All strains described in the above section are detailed in Table 1.

**Table 1:** Description of Ap7M, reversion, positive and functional mutant strains sequences. All viral strains produced and used in this study are detailed above. *The Amino Acid (AA) position is given with reference to the beginning of mature protein sequence (*ie*. AA position 107 in prM, written *prM* 107, corresponds to AA position 228 within the precursor polyprotein). A detailed map of protein positions within the YFV polyprotein sequence is provided in the supplementary Table S4. Synonymous mutations are indicated by grey, italic, lowercase letters. For each strain, the AA type encountered at a given position is indicated by the corresponding AA single letter code. The nucleotide substitutions corresponding to both synonymous and non-synonymous mutations are listed in the supplementary Table S5.

| Mutations introduced in *Asibi* strain AY640589 |  |  |  |  |  |  |  |  |  |  |  |  |  |  |  |
| --- | --- | --- | --- | --- | --- | --- | --- | --- | --- | --- | --- | --- | --- | --- | --- |
| Position<br>in<br>protein* | *Asibi* | *Ap7M* | Reversion mutant |  |  |  |  |  | Positive mutant |  |  |  | Functional mutant |  |  |
|  |  |  | E/H27Q | E/G28D | E/A155D | E/R323K | E/R331K | NS2A/A48T | E/Q27H | E/D155A | E/Q27H-D155A | E/Q27H-D28G-D155A | E/Q27A | E/Q27N | E/T154A |
| *prM*-107 | K | *k* | *k* | *k* | *k* | *k* | *k* | *k* | K | K | K | K | *k* | *k* | *k* |
| *prM*-136 | L | *l* | *l* | *l* | *l* | *l* | *l* | *l* | L | L | L | L | *l* | *l* | *l* |
| *E*-9 | R | *s* | *s* | *s* | *s* | *s* | *s* | *s* | R | R | R | R | *s* | *s* | *s* |
| *E*-27 | Q | H | Q | H | H | H | H | H | H | Q | H | H | A | N | H |
| *E*-28 | D | G | G | D | G | G | G | G | D | D | D | G | G | G | G |
| *E*-155 | D | A | A | A | D | A | A | A | D | A | A | A | A | A | A |
| *E*-323 | K | R | R | R | R | K | R | R | K | K | K | K | R | R | R |
| *E*-331 | K | R | R | R | R | R | K | R | K | K | K | K | R | R | R |
| *NS1*-109 | D | *d* | *d* | *d* | *d* | *d* | *d* | *d* | D | D | D | D | *d* | *d* | *d* |
| *NS2A*-48 | T | A | A | A | A | A | A | T | T | T | T | T | A | A | A |

### Recovery of infectious viruses and stock production

For producing each of the different YFV variants reported in this study, 3 to 4 overlapping DNA fragments were synthesized *de novo* (Genscript) and amplified by High Fidelity PCR using the Platinum PCR SuperMix High Fidelity kit (Life Technologies) and specific sets of primers. The first and last subgenomic fragments were flanked respectively at 5’ and 3’ termini by the human cytomegalovirus promoter (pCMV) and the hepatitis delta ribozyme followed by the simian virus 40 polyadenylation signal (HDR/SV40pA).

In the case of positive mutant strains, High Fidelity PCR amplification and fusion of subfragments FI.1.1 and FI.1.2 allowed the introduction of the required mutations into the subgenomic DNA fragment FI.1 bis (see Fig 1). After amplification, all DNA fragments were purified using Monarch PCR & DNA Cleanup kit 5μg (BioLabs) according to the manufacturer’s instructions.

Details regarding subgenomic DNA fragments and sets of primers used for the amplification and fusion steps are available in the supplementary Table S1. The different production schemes and subgenomic fragments combinations are described in Fig 1. Sequences of pCMV and HDR/SV40pA were described by Aubry and colleagues (38).

A final amount of 1μg of DNA (equimolar mix of subgenomic cDNA fragments) was transfected using Lipofectamine 3000 (Life Technologies) in a 25cm^2^ culture flask of subconfluent cells containing 1 mL of culture medium without antibiotics. Cell supernatants were harvested at 9 days post-transfection, aliquoted and stored at -80 °C. Each virus was then passaged two times onto BHK21 cells. Clarified cell supernatants from the second passage (virus stocks) were used to perform viral RNA quantification, TCID50 assays and whole-genome sequencing (see corresponding sections).

### Nucleic acids extraction

Samples (liver homogenate or cell culture supernatant) were extracted using either EZ1 Biorobot (EZ1 Virus Mini kit v2) or the QiaCube HT device and CadorPathogen kit both from Qiagen. Inactivation was performed using either 200μL of AVL buffer (EZ1) or 100μL of VXL buffer and 100 μL HBSS (Qiacube) according to the manufacturer’s instructions.

### Quantitative real-time RT-PCR assays

All quantitative real-time PCR (qRT-PCR) assays were performed using the EXPRESS SuperScript kit for One-Step qRT-PCR (Invitrogen). Primers and probe sequences are detailed in the supplementary Table S2.

The amount of viral RNA was calculated from standard curves using a synthetic RNA transcript (concentrations of 10^7^, 10^6^, 10^5^ and 10^4^ copies/μL). Results were normalized using amplification (qRT-PCR) of the housekeeping gene actin (as described by Piorkowski and colleagues (39)). The standard curves generated for all the YFV-specific assays had coefficient of determination values (R2) >0.98 and amplification efficiencies were between 93% and 100%.

### *In vivo* experiments

Three-week-old female Syrian Golden hamsters were inoculated intra-peritoneally with 5.10^5^ TCID50 of virus in a final volume of 100μL of HBSS. In all experiments, a control group of 2 hamsters was kept uninfected. The clinical course of the viral infection was monitored by following (*i*) the clinical manifestation of the disease (brittle fur, dehydration, prostration and lethargy), (*ii*) weight evolution and (*iii*) death. Weight was expressed as a normalized percentage of initial weight (%IW) and calculated as follows:

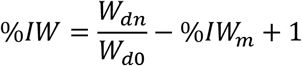

(*W_dn_*: weight at day n; *W_d0_*: weight at the day of the inoculation or challenge; *IW_n_*. mean of the %*IW* for control hamsters).

Liver samples were obtained from euthanized hamsters, grounded and treated with proteinase K before nucleic acid extraction using either the EZ1 Biorobot or the QiaCube HT device (see corresponding section).

### Whole and partial genome Next-Generation Sequencing

Both complete and partial coding nucleotide sequences were determined using next-generation sequencing methods: overlapping amplicons spanning either the complete genome sequence or the first part of the genome were produced from the extracted RNA (see corresponding section) using the SuperScript^®^ III One-Step RT-PCR System with Platinum®Taq High Fidelity kit (Invitrogen) and specific primers (detailed in the supplementary Table S3). Sequencing was performed using the PGM Ion torrent technology (Thermo Fisher Scientific) following the manufacturer’s instructions. Consensus sequence determination was done using CLC genomics workbench software (CLC bio-Qiagen). Substitutions with a frequency higher than 5% were considered for the analysis of intra-population genetic diversity and major/minor variants identification (minor variants: variants frequency >5% and <=75%; major variants: variants frequency >75%).

### Statistical analysis

Viral load comparisons were achieved using Wilcoxon rank-sum test and Kaplan-Meier survival analysis, using Mandel-Cox’s Logrank tests. Both analysis were performed using R software (40). P-values below 0.05 were considered as significant.

## RESULTS

### Reproduction of a lethal, viscerotropic infection in young female hamsters (*Mesocricetus Auratus*) using YFV strain *Asibi/hamster p7 Marseille* (Ap7M)

#### Strain design

In 2003, Mac Arthur and colleagues described a YFV hamster viscerotropic strain (*Asibi*/hamster p7) that was obtained after 7 serial passages of the *Asibi* (*Asibi*/hamster p0) strain in hamsters, (*e.g*. using the virus recovered from the liver of an infected hamster for subsequent infection) (15). The parental *Asibi*/hamster p0 strain is avirulent in hamsters, causing a limited viremia with no clinical manifestations whereas the derived *Asibi*/hamster p7 strain induces an important viremia (peaking at 3 days post-infection (pi)) along with a severe illness and death (mortality rate: 100%) in 3-week-old female hamsters.

Based on the reference strain *Asibi* (AN: AY640589), we designed the *Ap7M* virus sequence (see corresponding section in materials and methods). This strain includes 10 out of the 11 nucleotide changes identified in the *Asibi*/hamster p7 strain (15) in the coding region corresponding to proteins C to NS2A (6 non-synonymous, 5 synonymous). A comprehensive list of the codon changes in the open reading frame of the *Ap7M* strain with respect to the parental *Asibi* strain is displayed in Table 1. In addition, Table 1 details the codon changes for all the other mutants used in the current study.

The *Ap7M* virus was recovered after amplification and transfection of the 3 corresponding subgenomic amplicons in BHK21 cells according to the Infectious Subgenomic Amplicon (ISA) method (38), as detailed in Fig 1. The viral culture supernatant was then passaged twice onto BHK21 cells before storage, next-generation sequencing and titration of infectious particles.

We could not obtain the original strain *Asibi*/hamster p7 (*Ap7*), but the next passage on hamster livers (strain *Asibi*/hamster p8 (*Ap8*), which harbours a similar phenotype) was kindly provided by Dr Alan Barrett and Dr Patricia Aguilar (World Reference Center for Emerging Viruses and Arboviruses, University of Texas Medical Branch). *Ap8* virus was recovered from lyophilisate (resuspension in 500μL of HBSS) and passaged once in BHK21 cells before being stored, sequenced and titrated. It was used as a positive control for infection experiments. Strains *Asibi, Ap7* and *Ap8* are described in Fig 2. For all the viruses described in this study, viral titres are shown in Fig 3.

**Figure 3.**
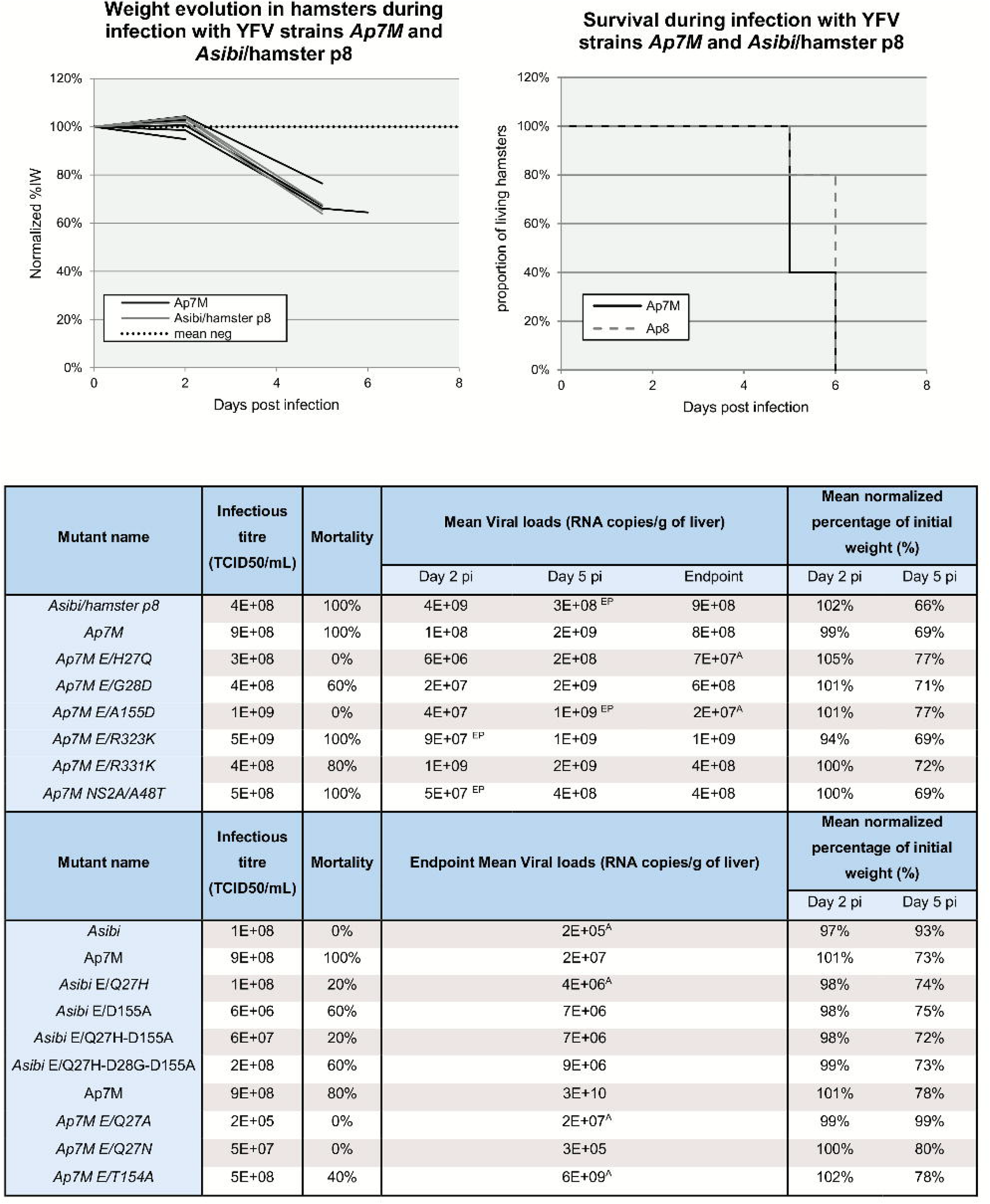
Ap7M and mutant strains in vitro and in vivo phenotypes. In the top left-hand corner, weight evolution in hamsters during infection with YFV strains *Ap7M* and *Asibi*/hamster p8 is detailed. Weight evolution is given for the 5 and 4 hamsters that were used for the evaluation of the mortality rate for *Ap7M* and *Ap8* strain, respectively. For the hamsters that were used as negative controls, weight evolution is expressed as a mean of % of initial weight. Hamsters infected with *Ap7M* strain are represented with black lines, those infected with *Ap8* strain, with grey lines and the mean for negative control, with a dotted line. In the top right-hand corner, the survival during infection with YFV strains *Ap7M* and *Asibi*/hamster p8 is illustrated. The evolution of the proportion of living hamsters is shown by a black line for *Ap7M* and by a grey dashed line for *Ap8*. It was calculated using 5 and 4 hamsters for *Ap7M* and *Ap8*, respectively. *In vitro* infectious titres, mortality rates, mean viral loads at 2, 5 dpi and 8 dpi/pm as well as mean normalized percentage of initial weight means are given for all viral strains described in this study. Viral loads, normalized percentage of initial weight and infectious titre evaluations are detailed in the corresponding paragraphs within the materials and methods section. Significant difference between viral loads at 2, 5 dpi and viral loads in endpoint (wilcoxon rank-sum test p-value<0.05) was indicated with a “EP” superscript next to the corresponding viral loads’ mean. Significant viral loads difference observed between Ap7M and other viruses at 2, 5 dpi or in endpoint (wilcoxon rank-sum test p-value<0.05) was indicated using a “A” superscript next to the corresponding viral loads’ mean.

**Figure 4.**
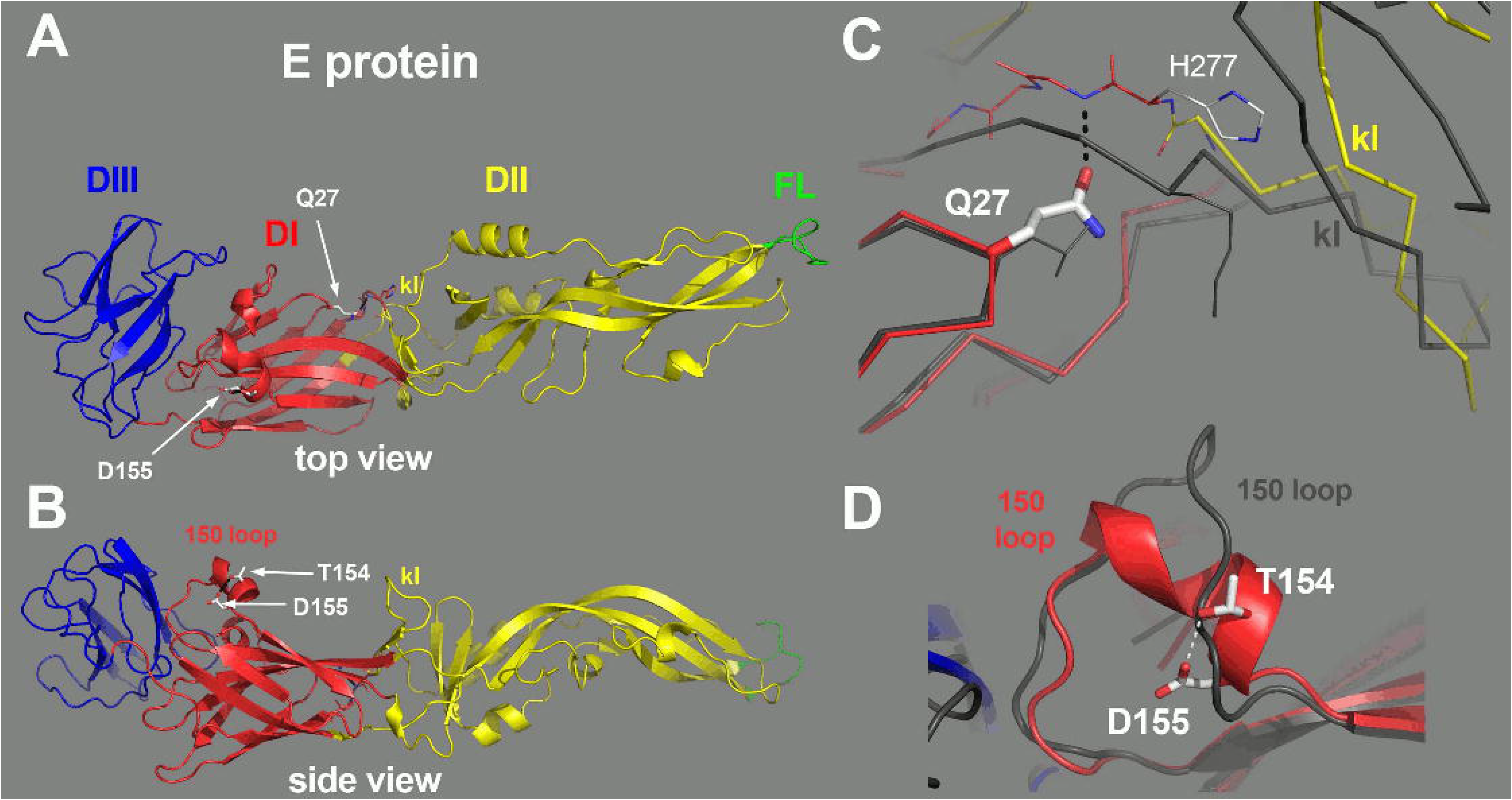
Location of the Q27H and D155A mutation on the structure of YFV E protein. The E protein is colored by domains, with domains I, II and III in red, yellow and blue, respectively, and the fusion loop highlighted in green. The “kl hairpin”, at the interface between domains I and II, is indicated. A) The E protein as viewed from the external surface of the virion, with the various domains labelled and the location of the mutations indicated by a white arrow. B) Side view, with the membrane below and the external side on top. The residues 154 and 155 are indicated, within a two-turn alpha helix in the 150 loop. C) Close-up view of the interactions of Glutamine 27 (Gln27). The side chain of Gln27 is in an extended conformation, making an interaction with the polypeptide chain at the end of the kl hairpin (labelled). For comparison, the E protein from TBEV was superposed on domain I, and is represented by a dark grey trace. Although the main chains superpose on domain I, in particular at the level of Q27, the *kl* hairpin is maintained away with respect to the TBEV *kl* hairpin (also labelled, in grey) by virtue of the interactions made by the Q27 side chain. This feature suggest that mutation of this amino acid will affect the conformation of this sensitive zone. D) The 150 loop of YFV E protein. For comparison, the TBEV E protein is shown superposed, in grey. The interaction between the D155 and T154 side chains is indicated, potentially stabilizing the alpha-helical conformation of this loop.

#### Infection

Two groups of 12 three-week-old female hamsters were inoculated intra-peritoneally with 5.10^5^ TCID50 of virus (either *Ap7M* or *Ap8* strain). A control group of 2 uninfected hamsters was maintained for weight monitoring. Clinical follow-up included (*i*) the clinical manifestations of the disease (brittle fur, dehydration, prostration and lethargy), (*ii*) body weight (weight evolution was expressed as a normalized percentage of the initial weight (%IW)) and (*iii*) death. Respectively 3 and 4 hamsters were euthanized at days 2 and 5 post-infection (dpi) to conduct virology investigations from liver samples, whilst the 5 others were kept for evaluating mortality rate (endpoint in case of survival: 8 dpi). Virology follow-up was achieved by performing qRT-PCR and next generation sequencing on RNA extracted from liver homogenates. All details regarding the *in vitro* and *in vivo* phenotypes of strains *Ap7M* and *Ap8* are available in Fig 3.

The integrity of the genome of both viruses was checked using Next-Generation Sequencing (NGS) methods. No additional mutation was identified in *Ap7M* strain sequence whereas, for the *Asibi*/hamster p8 strain, the E/D28G non-synonymous mutation was found to be reverted (*i.e*., similar to the sequence of the *Asibi* strain) for both the original *Ap8* lyophilisate and the stock produced in BHK21 cells (see Fig 2).

Nearly all hamsters inoculated with Ap7M and Ap8 strains developed outward signs of illness such as brittle fur, prostration, dehydration and lethargy. One hamster from the *Ap8* infected group did not show any sign of illness, both its liver and blood were tested negative for YFV genome at 8 dpi and it was excluded from analysis. Maximal mortality rates (100%) were observed for both *Ap7M* and *Ap8* strains (see Fig 3), similar to previous observations by McArthur *et al*. for *Ap7* strain infection in hamsters (15). Clinical signs of the disease appeared as early as 4 dpi and all animals died within two days after onset of the symptoms. No weight loss was observed at 2 dpi (means of %IW of 99% and 102% for *Ap7M* and *Ap8* virus, respectively). In contrast, severe weight loss was recorded at 5 dpi in both groups (means of %IW of 69% and 66%, respectively).

All livers were found to be YFV-positive by qRT-PCR in both infected groups. Viral RNA loads in the liver samples ranged between 1.10^8^ and 4.10^9^ RNA copies per gram of liver. There was no significant difference observed between the viral yields at 2 and 5 dpi and endpoint viral loads in the Ap7M-infected group. In the *Ap8* group, a significant increase was observed between day 5 pi and endpoint viral loads (Wilcoxon rank-sum test (WRST), p=0.029). For both groups, viral loads’ evolution in the liver suggests a peak period between 3 and 4 dpi.

Next Generation Sequencing (NGS) was performed from liver homogenates recovered from Ap7M-infected hamsters. Sequencing of the liver samples was achieved on the structural proteins’ coding region (ORF nucleotide positions 1-3879) and performed for the five hamsters used for mortality rate evaluation. For intra-population genetic diversity analysis, a variant was regarded as major when the corresponding nucleotide proportion (CNP) was over 75%. All sequencing results are shown in Table 2. In the case of the *Ap7M* strain, no change in the viral sequence was detected after propagation in hamsters. The *in vivo* phenotype observed for this strain in hamsters is thus attributable to the 10 mutations introduced into the *Asibi* strain genome.

**Table 2:**
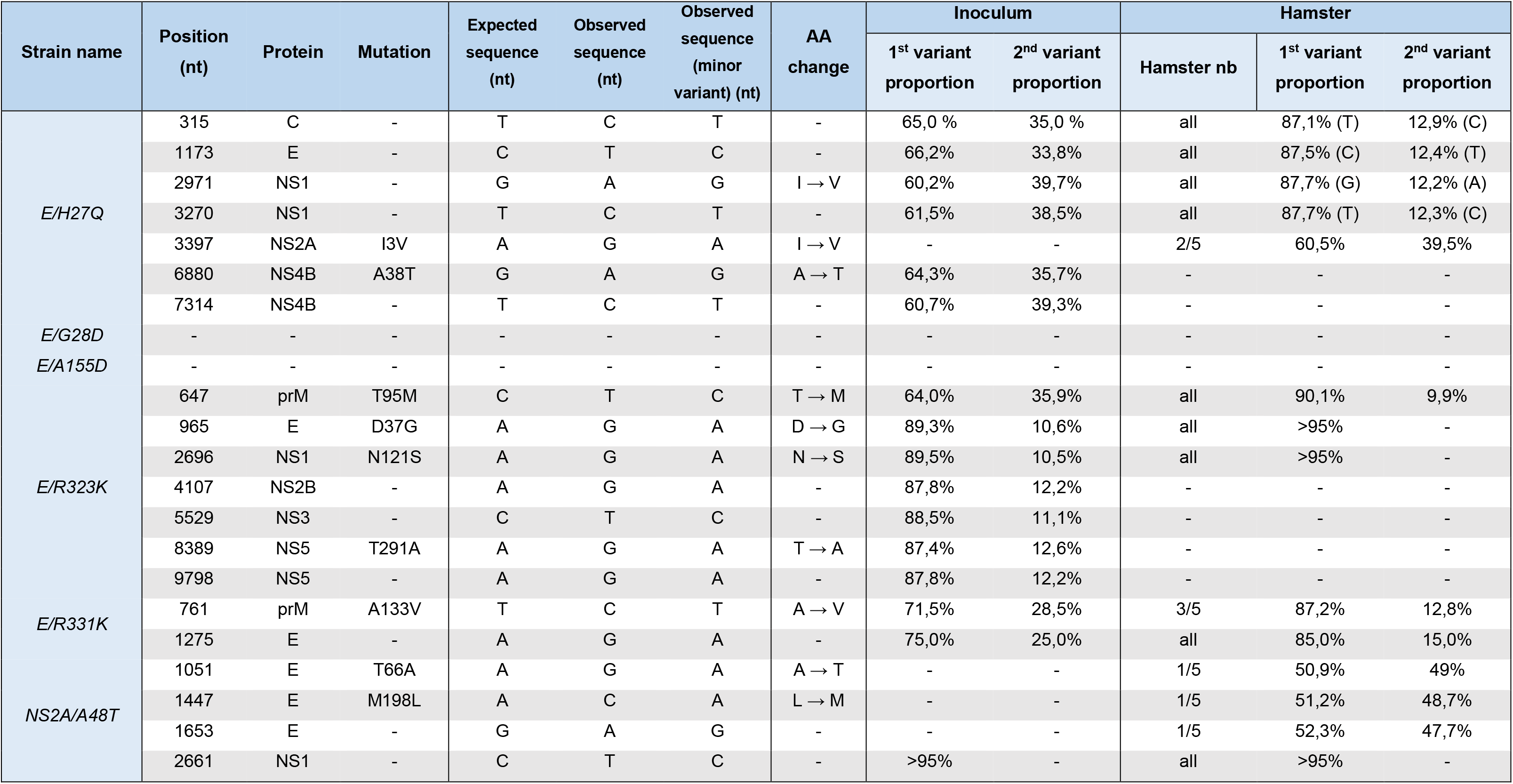
Sequence and intra-population diversity from viral culture supernatant and hamster liver samples for mutant strains Ap7M E/H27Q, E/G28D, E/A155D, E/R323K, E/R331K and NS2A/A48T. For each reversion mutant, the changes found either in the complete coding sequence from the viral stock or in partial sequences from infected hamster liver homogenates are detailed above. For mutations identified in more than one sequence from hamster liver, 1^st^ and 2^nd^ variants proportions are given as means. Expected sequence refers to that of plasmids that served as templates for subgenomic fragment amplification while producing viruses thanks to the ISA method.

### Evaluation of the impact on virulence of the reversion of *Ap7M* non-synonymous mutations in the E and NS2A genes

#### Strain design

In total, 6 deletion mutants were designed by selectively deleting (*i.e*. reverting to *Asibi* strain sequence) each of the non-synonymous mutations present in the E (5 mutations) and NS2A (1 mutation) genes of *Ap7M*. Details regarding the sequences of the reversion mutants *Ap7M* E/H27Q (E/H27Q), *Ap7M* E/G28D (E/G28D), *Ap7M* E/A155D (E/A155D), *Ap7M* E/R323K (E/R323K), *Ap7M* E/R331K (E/R331K) and *Ap7M* NS2A/A48T (NS2A/A48T) are provided in Table 1. Each of the deletion mutants was produced using 4 subgenomic amplicons that were amplified and transfected in BHK21 cells (see Fig 1). After transfection, all viruses were passaged, stored, sequenced and titrated as described for *Ap7M* virus.

#### Infection design

Infection, clinical and virology follow-up as well as NGS were achieved as described for strain *Ap7M*. Nine hamsters were inoculated with mutant strain E/G28D and 2 hamsters were used for virology investigations at 2 and 5 dpi. All details regarding the *in vitro* and *in vivo* phenotypes of reversion mutant strains are available in Fig 3.

#### Infection by strains E/H27Q, E/G28D and E/A155D

Both complete coding sequences obtained from cell culture supernatants (viral stocks) of mutant strains E/A155D and E/H27Q included only the desired reversion. The E/A155D sequence was strictly identical to the sequence designed *in silico*, but the E/H27Q sequence had six extra mutations (2 non-synonymous and 4 synonymous), although all of them were associated with minor variants (CNPs between 60 and 65%) (Table 2).

No death was observed in groups infected with E/H27Q and E/A155D reversion mutants. This dramatic change in mortality rate came along with higher means of %IW at 5 dpi (77% for both groups, respectively) and lower viral loads’ means at 2 dpi (6.10^6^ and 4.10^7^ for E/H27Q and E/A155D-infected groups, respectively) and at endpoint (7.10^7^ and 2.10^7^ for E/H27Q and E/A155D-infected groups, respectively, with WRST p=0.007937 for both viruses). Analysis of the partial sequences obtained from liver homogenates of E/A155D-infected hamsters showed no additional change to the viral sequence. Besides, mutations associated to minor variants in E/H27Q viral stock sequence were found to be reverted when sequencing the liver samples. An additional non-synonymous change (I3V) was identified in the NS2A coding region of the sequences from two liver samples (see Table 2) but was associated to a minor variant in both cases (CNPs of 59,2% and 61,8%, respectively). Hence, the change in the *in vivo* phenotype observed for reversion mutants E/H27Q and E/A155D is undoubtedly imputable to the reversion of mutations E/Q27H and E/D155A, respectively.

In the case of mutant E/G28D, the complete coding sequence obtained from the viral stock included only the desired reversion. In the E/G28D-infected group, the mortality rate was decreased (60%) and a slight reduction of the viral load was observed at 2 dpi (mean: 2.10^7^). However, means of %IW at days 2 (101%) and 5 pi (71%) and mean viral loads at 5 dpi (mean: 2.10^9^) and at endpoint (6.10^8^) were similar to those observed in the Ap7M-infected group. The sequences obtained from infected hamster livers homogenates were similar to those of the inoculum (viral stock). In consequence, the E/D28G mutation reversion can account for the mild changes in the *in vivo* phenotype that were observed.

#### Infection by strains E/R323K, E/R331K and NS2A/A48T

In the case of E/R323K, E/R331K and NS2A/A48T mutant strains, mutations associated to major variants were identified in complete sequences from the viral stocks. One synonymous mutation was identified in the NS1 coding region of NS2A/A48T. Six mutations were detected in the sequence of E/R323K mutant strain (3 NS and 3 S). Amongst the 3 NS changes, only one was located in the E coding region (D37G). In the sequence from mutant E/R331K viral stock, two mutations were observed (1 NS and 1 S) including a NS change located in the prM region (A133V). Only the synonymous change was associated to a major variant.

In groups infected with mutants E/R323K, E/R331K and NS2A/A48T both clinical and virological evolutions were similar to those observed in the model infection (*Ap7M* strain). Clinical signs of illness appeared between 4 and 5 dpi, shortly followed by death with mortality rates between 80 and 100%. As observed for Ap7M-infected group, means of %IW were close to 100% at 2 dpi (94, 100 and 100% for E/R323K, E/R331K and NS2A/A48T-infected groups, respectively) and decreased dramatically at day 5 pi (69, 72 and 69%, respectively). Mean viral loads ranged from 5.10^7^ to 1.10^9^ at 2 dpi, with a 1-log increase between day 2 and 5 pi, and peaked around 3 dpi.

In the case of strain NS2A/A48T, the undesired NS1 synonymous mutation was found again in the partial sequences from liver samples. Three additional changes (2 NS and 1 S) were detected in the E coding region (T66A, M198L) from one of the liver homogenates but none of them was associated to a major variant (CNPs between 51% and 52%). The preservation of an *in vivo* phenotype similar to that of *Ap7M* and the absence of NS mutations associated to major variants suggest a subordinate role of mutation NS2A/T48A in *Ap7M* strain virulence.

Regarding strain E/R331K, both undesired mutations detected in the viral stock sequence were also identified as major variants in the partial sequences obtained from hamster livers (CNPs >80%). For strain E/R323K, the envelope non-synonymous change D37G was identified as a major variant (CNPs >90%) in all partial sequences from hamster livers. Although for both strains, the *in vivo* phenotype was close to that of *Ap7M*, owing to the presence of additional NS mutations in the prM and E genes of E/R323K and E/R331K strain, respectively, no definitive conclusion can be drawn regarding the phenotypic impact of E/K323R and E/K331R mutations *in vivo*.

### Evaluation of the phenotypic impact of the introduction of *Ap7M* mutations E/Q27H, E/D28G and E/D155A in the genome of YFV strain *Asibi*

#### Strain design

A clear phenotypic impact was evidenced for the reversion of mutations E/Q27H, E/D28G and E/D155A. Based on these results, 4 “positive” mutants were designed by selectively introducing one or more of these mutations into the sequence of YFV strain *Asibi: Asibi* E/Q27H, *Asibi* E/D155A, *Asibi* E/Q27H-D155A and *Asibi* E/Q27H-D28G-D155A. These mutants are described in Table 1. As for the reversion mutants, all *Asibi* “positive” mutants were produced by amplifying and transfecting 4 subgenomic amplicons in BHK21 cells. Additional steps of amplification and fusion were performed to introduce the required mutations into the first DNA fragment of YFV *Asibi* strain FI.1bis (see Fig 1). All viruses were subsequently passed, stored, sequenced and titrated, as described above.

#### Infection design

Infection, clinical and virology follow-up as well as NGS were achieved as described for strain *Ap7M*. Groups were limited to 5 hamsters that were used for mortality rate evaluation and virus detection in the liver either at 8 dpi or *post-mortem*. As controls to this experiment, two groups of 5 hamsters were infected with YFV strains *Ap7M* and *Asibi,* respectively. The *in vitro* and *in vivo* phenotypes of all positive mutant strains are described in Fig 3 and NGS results are shown in Table 3.

**Table 3:**
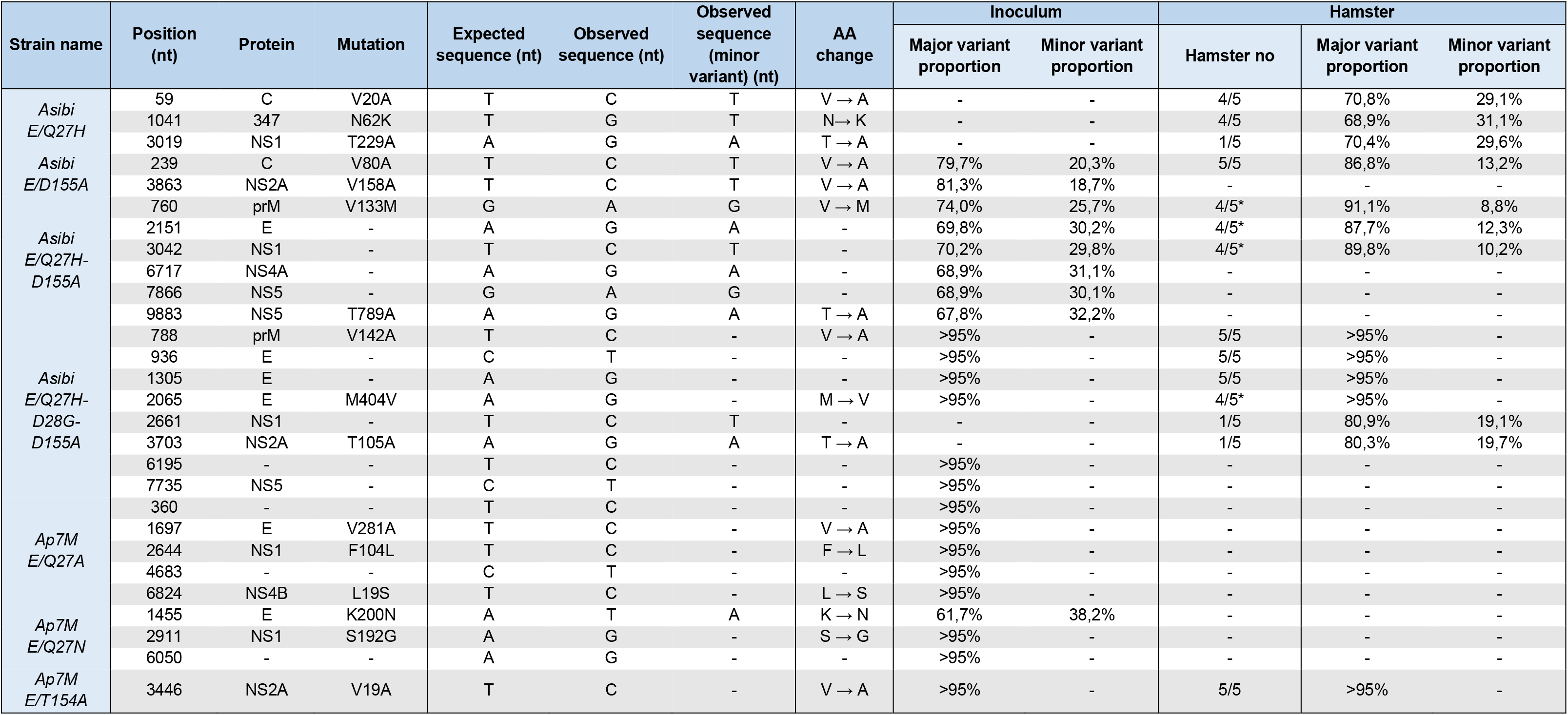
Sequence and intra-population diversity from viral culture supernatant and hamster liver samples for mutant strains Asibi E/Q27H, E/D155A, E/Q27H-D155A, E/Q27E1-D28G-D155A, Ap7m E/Q27A, E/Q27N, E/T154A. For each reversion mutant, the changes found either in the complete coding sequence from the viral stock or in partial sequences from infected hamster liver homogenates are detailed above. For mutations identified in more than one sequence from hamster liver, 1^st^ and 2^nd^ variant proportions are given as means unless no 2^nd^ variant was detected (>95%). Expected sequence refers either to that of plasmids or to that of primers (used for mutation introduction) that served for subgenomic fragment amplification. For *Asibi* E/Q27H-D155A and *Asibi* E/Q27H-D28G-D155A mutants, some mutations were found to be reverted in one out of the five hamsters as indicated by the star (*). In this case, the 1^st^ and 2^nd^ variants proportions were not included for means’ calculation

Results obtained with the Ap7M-infected group were comparable to those previously observed. The mortality rate was maximal (100%), with clinical signs of illness appearing as early as 4 dpi and a severe weight loss observed between 2 and 5 dpi (mean of %IW: 101 and 73%, respectively). Viral loads were slightly inferior to those previously reported in Ap7M-infected hamsters (mean: 2.10^7^). The unusually long disease of one of the hamsters in Ap7M-infected group led to the extension of the experiment to 9 dpi.

No death and nearly no disease was observed in the Asibi-infected group. The clinical signs of illness were limited to a mild loss of weight at 5 dpi (mean of %IW: 93%) and endpoint viral loads were 2 logs lower than those observed in the model infection (mean: 2.10^5^, WRST p=0.007937).

All groups infected with *Asibi* “positive” mutant strains showed signs of severe viscerotropic disease.

#### Infection by strain Asibi E/D155A

Two non-synonymous changes were detected in the sequence from the viral stock of mutant strain *Asibi* E/D155A in the C (V80A) and NS2A (V158A) proteins and were both identified as major variants (CNPs of 79,7 and 81,3%, respectively). A high mortality rate (60%) was observed in E/D155A-infected group and both clinical and virological evolutions were similar to those observed in hamsters infected with *Ap7M*. Mean %IW decreased from 98% at 2 dpi to 75% at 5 dpi and mean endpoint viral loads were close to those observed in the Ap7M infection (7.10^6^). All hamsters showed dehydration signs and brittle fur, and prostration was recorded for most of them (60%, data not shown). When sequencing liver samples from infected hamsters, the C protein NS mutation was still present in all samples. Since no additional mutation was observed in the E protein gene, it is probable that the phenotypic change observed is attributable to the E/D155A mutation.

#### Infection by strain Asibi E/Q27H

The complete coding sequence obtained from the supernatant of cells infected with the Asibi E/Q27H strain was similar to the *in silico* designed sequence. A moderate mortality rate (20%) was recorded amongst E/Q27H-infected hamsters. Important weight loss was observed between 2 dpi (mean of %IW: 98%) and 5 dpi (mean of %IW: 74%) but the endpoint viral loads mean was significantly lower than that of the *Ap7M* infection (4.10^6^). Whilst all hamsters showed dehydration signs and brittle fur, prostration was observed in only one of them (data not shown). All non-synonymous mutations identified in the 4 partial sequences obtained from infected hamster liver homogenates (1 partial sequence could not be obtained) corresponded to minor variants (CNPs between 69% and 71%). Altogether, the moderate phenotypic change observed in positive mutant strain E/Q27H, can be attributed to mutation E/Q27H.

#### Infection by strain Asibi E/Q27H-D155A

In the case of strain *Asibi* E/Q27H-D155A, 6 mutations were detected in the complete coding sequence of the virus, including 2 NS changes located in proteins prM (V133M) and NS5 (T789A), all associated with minor variants (CNPs between 68 and 74%). In E/Q27H-D155A infected groups, the mortality was moderate (20%), accompanied by an important weight loss between 2 and 5 dpi (from a mean of %IW of 98% to 72%), endpoint viral loads close to that observed in the model infection (mean: 7.10^6^) and additional clinical signs of illness limited to dehydration signs and brittle fur (data not shown). The NS prM change V133M, was found in the partial sequences from 4 of the infected hamsters and corresponded to major variants (CNPs of 87.7 and 91.1%). It was found to be reverted in the last one. This experimental set globally confirms that mutations at positions E/Q27H and E/D155A modulate virulence in hamsters.

#### Infection by strain Asibi E/Q27H-D28G-D155A

The sequence of *Asibi* E/Q27H-D28G-D155A mutant showed 6 additional mutations (2 NS and 4 S), all associated to major variants (CNPs>95%), notably affecting prM (V142A) and E (M404V) proteins. A high mortality rate (60%), important weight loss between 2 and 5 dpi (from a mean of %IW of 99% to 73%) and endpoint viral loads close to that observed in the model infection (mean: 9.10^6^) were observed. Dehydration signs, brittle fur and prostration were reported in nearly all cases (80%, data not shown). The envelope M404V change, was detected and associated to major variants in the majority of the sequences (4/5) but was found to be reverted in the last one. Interestingly, two additional mutations were identified in this sequence. They both corresponded to major variants (CNPs of 80.3 and 80.9%) and included a NS change in the NS2A protein (T105A). The corresponding hamster died of viral infection so no specific phenotype could be attributed to those mutations. Those results globally confirm that variants at positions E/Q27H, E/D28G and E/D155A can strongly modify virulence in hamsters.

### Location and structural impact of mutations Q27H, D28G and D155A in the E protein of YFV strain *Asibi*

The amino acid substitutions Q27H and D155A were identified as sufficient to induce a pathogenic phenotype in hamsters for the *Asibi* strain (see above). In the structure of the YFV E protein (*Asibi* strain, pdb accession code 6EPK), both residues Q27 and D155 are in domain I and their side chains are involved in polar interactions that could affect the conformation of the protein. Residues Q27 and D28 are at a β-turn connecting strand B0 and C0 of domain I. The Q27 side chain accepts a hydrogen bond from the main chain amide of G278 at the end of the *kl* hairpin of domain II. This interaction therefore imposes constraints in the conformation of the polypeptide chain in the hinge area between domains I and II, which has been reported earlier to affect flavivirus virulence (41). Modeling a Histidine residue at this position resulted in conformers that could not make the same interactions as Q27, because the imidazole ring in the side chain projected out differently. Mutation of the Q27 residue to an asparagine (which would not be long enough to make the same interactions as Q27) or to an alanine should affect the interactions observed between Q27 and the main chain in a way analogous to that observed with H27 residue.

D155 is in the “150 loop” of domain I, a variable segment across the flavivirus E proteins which is exposed at the surface of the virion; the E proteins from many flaviviruses carry a glycosylation site in this loop (42-45). Although there is no N-linked glycan attached to YFV E protein, the structure shows that D155 is in a short alpha helix, and its side chain makes a hydrogen bond with that of the preceding residue, T154, which is likely to stabilize the conformation of the helix. If the disruption of this interaction is important for the virulent phenotype of *Ap7M*, mutation of T154 residue to alanine would also disrupt this interaction, with an effect similar to that of the D155A mutation and thus, induce virulence in hamsters.

### Phenotypic impact of the substitution of mutation Q27H with Q27A or Q27N and of mutation D155A with T154A into the E protein sequence of YFV strain *Ap7M*

#### Strain design

To test the potential role of the interactions observed by G27 and D154 in the Asibi strain, the Q27H and D155A, “functional” mutants were designed by substituting those two mutations with the potentially functionally similar mutations discussed above.

In the E/Q27A and E/Q27N functional mutants, the E/Q27H histidine (in the *Ap7M* strain) was replaced by an alanine or an asparagine residue, respectively. In the E/T154A functional mutant, the E/D155A alanine (in *Ap7M* strain) was reverted to the original aspartic acid (in *Asibi* strain) whilst the amino acid in position E/T154A was changed from a threonine to an alanine (see Table 1).

All functional mutants were produced using 4 subgenomic amplicons that were amplified from the corresponding plasmids and transfected in BHK21 cells (see Fig 1). Following transfection, the viruses were passaged, stored, sequenced and titrated similarly to the other viruses.

#### Infection design

Infection, clinical and virological follow-up as well as NGS were achieved as described for the positive mutant strains. As a control for this experiment, a group of 5 hamsters was infected with YFV strain *Ap7M*. Next-generation sequencing on the liver homogenates recovered from infected hamsters was only performed for *Ap7M* E/T154A virus (results shown in Table 3). All the details regarding means of %IW, viral loads’ means and mortality rates are available in Fig 3.

All hamsters from the Ap7M-infected group were affected by a severe viscerotropic disease, with the first outward signs of illness appearing from 4 dpi. Death was observed as soon as 5 dpi and until 8 dpi, a delay accompanied by a mortality rate slightly lower than that previously observed (80%). Although mean of %IW at 5 dpi was higher than in the other experiments (78%), severe weight loss was subsequently observed at 6, 7 and 8 dpi (mean of %IW of 68, 61 and 55%, respectively) and high endpoint viral loads (mean: 3.10^10^) were recorded.

#### Infection by strain E/T154A

One additional NS mutation was detected in the sequence of strain *Ap7M* E/T154A in the region coding for the NS2A protein (V19A) and was associated to a major variant (CNPs >95%). The hamsters infected with E/T154A mutant strain nearly all showed signs of a severe viscerotropic disease (4 out of 5 individuals). Although the mortality rate was lower (40%), both clinical and virological evolutions were close to those observed in the group infected with strain *Ap7M*. Mean of %IW fell from 102% at 2 dpi to 78% at 5 dpi and decreased until 64% at 8 dpi. Viral loads were high and varied between 5.10^8^ and 3.10^10^, with no significant difference compared to *Ap7M* infection. The NS change V19A was identified as associated to a major variant in the partial sequences from all liver homogenates with no additional change. Since no additional mutation was observed in the E gene, it is very likely that the restoration of *Ap7M* virulence in hamsters is attributable to the E/T154A mutation.

#### Infection by strain E/Q27N

The sequence of mutant *Ap7M* E/Q27N included 3 additional mutations including 2 NS changes located in the E (K200N) and in the NS1 (S192G) coding regions. As the residue 202 in E protein is a serine, the K200N change introduces a new N-glycosylation site within the E protein. Only the NS1 mutation corresponded to a major variant. Although no death was observed in the group infected with E/Q27N mutant, all hamsters showed signs of illness, including an important loss of weight (mean of %IW of 80% at 5 dpi and between 74 and 75% from 6 dpi to 8 dpi). However, endpoint viral loads were significantly reduced compared to that observed with strain *Ap7M* (mean: 3.10^5^, WRST p=0.007937). Although a NS mutation was identified in the NS1 gene of mutant E/Q27N, there are grounds to believe that mutation E/Q27N did not allow the recovery of the *in vivo* phenotype of *Ap7M* and therefore, did not provide a functional equivalent to the E/Q27H mutation observed in *Ap7M* strain.

#### Infection by strain E/Q27A

In the sequence of the functional mutant strain *Ap7M* E/Q27A, 6 extra mutations were identified (3 NS and 2 S). NS changes affected E (V281A), NS1 (F104L) and NS4B (L19S) proteins and were all associated to major variants (CNPs>95%). Hamsters from the E/Q27A-infected group showed no clinical signs of disease and no mortality nor loss of weight was recorded in this group (minimum mean of %IW: 99%, reached at 5 dpi). In addition, endpoint viral loads were low, *i.e*. comprised between 0 and 1.10^6^ (WRST p=0.01116). The observed *in vivo* phenotype cannot be reliably associated to mutation E/Q27A since 3 additional NS mutations were identified in the mutant protein sequence, notably in the envelope gene (V281A).

## DISCUSSION

This study started with providing a proof of concept that the recently described ISA (Infectious Subgenomic Amplicons) reverse genetics method (38) could be used to produce genetically engineered flaviviruses with specific phenotypic properties. Here, we first produced an equivalent to the hamster-virulent strain Yellow Fever *Ap7* (15) by simply introducing a set of 10 mutations into a single subgenomic amplicon derived from the sequence of the *Asibi* strain. When inoculated to young female hamsters, the resulting strain, YF *Ap7M*, induced a lethal viscerotropic disease similar to that described for YF *Ap7* virus in terms of clinical signs of illness, weight evolution, viral loads in the liver and lethality (100%). We further exploited the ISA methodology by producing a set of mutant strains derived from either *Ap7M* or *Asibi* viruses and allowing to investigate for each of the non-synonymous mutations differentiating *Ap7M* from *Asibi* (alone or combined), the impact on the *in vivo* phenotype of the virus. The approach was classical, but it is noticeable that the absence of requirement for cloning a complete modified genome allowed the simple and convenient production of multiple mutants from subgenomic synthetic fragments in days, with a one hundred percent success in virus recovery. Then, the combination of the ISA reverse-genetics approach and next-generation sequencing proved to be critical to correctly associate the mutation(s) introduced to the *in vivo* phenotype of the mutants. Obviously, a process excluding exhaustive genetic data analysis would have led, in some cases, to inaccurate conclusions. Indeed, we detected undesired non-synonymous mutations, notably in the E protein coding region of several sequences retrieved from the hamster liver samples. These mutations polluted the interpretation however, our results indicate that some of the changes introduced within the E protein sequence have a significant impact on the virulence phenotype of the mutant strains, suggesting that the corresponding amino acid residues are involved in interactions that are relevant for the pathogenic features of yellow fever virus.

Two of the E protein variant residues turned out to be key components of the virulence mechanism in hamsters. The reversion in *Ap7M* of either E/Q27H or E/D155A mutations to the original *Asibi* residue, led to an important reduction of virulence: lethality was abolished, viral loads were reduced, and clinical signs of illness were, to a variable extent, alleviated. In addition, the introduction of the single D155A *Ap7M* mutation into the E protein of the YF strain *Asibi*, was sufficient to drastically modify its *in vivo* phenotype in hamsters towards both a greater replication efficiency (*i.e*. with higher viral loads) and virulence (*i.e*. with increased signs of illness and lethality). The impact of other *Ap7M* mutations could not be fully determined because production in cell cultures was associated with the appearance of undesired non-synonymous mutations.

The structure of YFV E protein (*Asibi* strain, PDB 6EPK) shows that the side chains of residues Q27 and D155 are involved in polar interactions that could impact the overall conformation of the protein. Using functional mutants, we tested the implication in YF *Ap7M* virulent phenotype of both modifications of the interactions between Q27 and the main chain (at position E278) and between T154 and D155 residues. We replaced H27 residue (in *Ap7M*) with the residues A27 and N27, which would not be able to make the same interactions as Q27, but failed to restore the *Ap7M in vivo* phenotype. In contrast, replacement of D155A with a T154A change within the sequence of the *Ap7M* sequence allowed the recovery of a hamster-virulent phenotype. This result provides a structural basis for the relationship between mutation E/D155A and the virulence in hamsters observed with the *Ap7M* virus.

The identification of residues in the yellow fever virus E protein carrying the virulence phenotype in hamsters is of utmost interest for the *in vivo* testing of antiviral molecules against YFV. It implies that the efficacy and the mechanism of action of antivirals targeting the non-structural proteins of the virus can be tested in a reproducible lethal hamster model (after providing the relevant mutations in the E gene), using genetic engineering of the target gene. Our study shows that this is possible using the genetic backbone of the *Asibi* strain but it remains to be established whether the determinants of virulence in this evolutionary lineage can be used to modulate virulence in other genotypes. An alignment of 169 YF envelope sequences indicates that E/Q27H and E/D155A mutations cannot be found in any other YF sequences with the exception of the *in vitro* adapted Asibi-LP-CDC HeLa p6 strain (15, 46), the virulence of which in hamsters has not been reported. Thus, although obtaining a hamster-virulent phenotype should not be taken for granted, there is a potential for rapidly modifying other YF strains in the prospect of *in vivo* studies. Of note, other yellow fever virus strains (Jimenez p0 and p10) have been reported to be virulent in hamsters (36). Unfortunately, the only available sequence (Jimenez p0, GenBank AN: AY540480) is partial and does not include the region of interest in the E gene. Further analysis of Jimenez p0 and p10 sequences should prove valuable to determine if these strains share some of the determinants of yellow fever virus virulence in hamsters described in our study.

Finally, the hamster-virulent strains described in this study are a tangible example of how a few mutations in the genome of flaviviruses can impact replicative fitness and virulence in an unusual host. Rodents have never been reported as hosts for YFV in nature and primary isolates do not grow well in hamsters (35). As described above, the *Asibi* strain causes only a limited and inapparent infection in hamsters. This work brings reasonable grounds to believe that, if flaviviruses were led by a conjunction of environmental and anthropic factors to cross species barrier, a very limited number of mutations could be enough for the virus to acquire an increased replicative fitness but also a virulent phenotype.

## ACKNOWLEDGEMENTS

We thank Grégory Moureau, Karine Barthélémy, Géraldine Piorkowski for technical assistance. We thank by Dr Alan Barett and Dr Patricia Aguilar (World Reference Center for Emerging Viruses and Arboviruses, University of Texas Medical Branch) for providing The *Asibi*/hamster p8 strain.

## CONFLICT OF INTEREST

The authors have declared that no competing interests exist.

## SUPPLEMENTARY MATERIALS

*Supplementary Protocol S1*. Protocols regarding cell culture (virus stock production and virus titration (TCID50)), PCR amplification and fusion (subgenomic amplicon production), transfection and virus stock production, quantitative real-time PCR (qRT-PCR) assays, sample collection as well as next-generation sequencing are detailed in this section.

*Supplementary Table S1. Subgenomic fragments and corresponding primers used for virus production using the ISA method*.

*Supplementary Table S2. Yellow Fever Virus and Actin-specifìc qRT-PCR systems*.

*Supplementary Table S3. Yellow Fever Virus-specific PCR systems used for High-Fidelity RT-PCR amplification*.

*Supplementary Table S4. Mature proteins positions within Yellow Fever Virus polyprotein sequence*. Obtained from UniProt database (www.uniprot.org, UniProt: the universal protein knowledgebase. Nucleic Acids Res. 45: D158-D169 (2017))

*Supplementary Table S5. Description ofAp7M, reversion, positive and functional mutant strains nucleotide sequences*. The Amino Acid (AA) position is given with reference to the beginning of mature protein sequence (ie. AA position 107 in prM, written prM107, corresponds to AA position 228 within the precursor polyprotein). A detailed map of protein positions within the YFV polyprotein sequence is provided as supplementary Table S4. Synonymous mutations are indicated by grey, italic, lowercase letters. For each strain, the AA type encountered at a given position is indicated by the corresponding AA single letter code.

